# Development and evaluation of a loop-mediated isothermal amplification (LAMP) assay for the detection of *Tomato brown rugose fruit virus* (ToBRFV)

**DOI:** 10.1101/2020.03.02.972885

**Authors:** Alian Sarkes, Heting Fu, David Feindel, Michael W. Harding, Jie Feng

## Abstract

*Tomato brown rugose fruit virus* (ToBRFV) is a member of *Tobamovirus* infecting tomato and pepper. Within North America, both the United States and Mexico consider ToBRFV to be a regulated pest. In Canada, the presence of ToBRFV has been reported, but an efficient diagnostic system has not yet been established. Here, we describe the development and assessment of a loop-mediated isothermal amplification (LAMP)-based assay to detect ToBRFV. The LAMP test was efficient and robust, and results could be obtained within 35 min with an available RNA sample. Amplification was possible when either water bath or oven were used to maintain the temperature at isothermal conditions (65°C), and results could be read by visual observation of colour change. Detection limit of the LAMP was eight target RNA molecules. Under the experimental conditions tested, LAMP was as sensitive as qPCR and 100 times more sensitive than the currently used rt-PCR. We recommend this sensitive, efficient LAMP protocol to be used for routine lab testing of ToBRFV.

## Introduction

Tomato (*Solanum lycopersicum* L.) is one of the most important vegetable crops in the world [1]. Canada is the main producer of greenhouse tomatoes in North America, with an annual yield of more than 250,000 t and a value of around $500 million (Agriculture & Agri-Food Canada, www.agr.gc.ca/pmc-cropprofiles). Diseases caused by viruses are one of the most critical factors affecting tomato production worldwide [2]. *Tomato brown rugose fruit virus* (ToBRFV) is a member of the *Tobamovirus* genus with a host range including tomato, pepper and some weeds such as nightshades [3]. It was first detected in Israel in 2014 [4] and later in countries in North America [5,6], Asia [7–9] and Europe [10,11]. Within North America, both the United States [12] and Mexico [13] consider ToBRFV to be a regulated pest. In Canada, ToBRFV was first identified in Ontario in 2019 (http://thegrower.org/news/tomato-brown-rugose-fruit-virus-identified-ontario). Since then the Alberta Plant Health Lab (APHL) began testing of tomato samples submitted from Alberta greenhouses, but no positive samples had been found at the time this report was submitted.

ToBRFV has a single-stranded positive-sense RNA of ~6,400 nucleotides (nt), with a typical *tobamovirus* genome organization that consists in four open reading frames (ORFs) encoding two replication-related proteins [7]. On tomato, ToBRFV causes symptoms including leaf interveinal yellowing and deformation, mosaic staining, young leaves deformation and necrosis, sepal necrosis and deformation, young fruits discoloration, deformation and necrosis [3]. Besides the significant damage to yield and fruit quality, ToBRFV is of special concern compared to other *tobamoviruses* because of its ability to overcome the *R* genes *Tm-2* and *L*, which are deployed in tomato and pepper varieties, respectively, for *tobamoviruses* resistance [4].

An effective disease management program is dependent on timely and proper identification to the causal agent of the disease. Enzyme-linked immunosorbent assay (ELISA), reverse transcriptase polymerase chain reaction (rt-PCR), next-generation sequencing (NGS) and transmission electron microscopy (TEM) have been used for ToBRFV detection [3]. In Canada, the Alberta Plant Health Lab, Alberta Agriculture and Forestry, had utilized the rt-PCR methods developed by Ling et al. [6] for diagnosis on suspected tomato samples. Recently, protocols employing real time rt-PCR for ToBRFV detection were developed and demonstrated to be highly specific and sensitive [14].

As an alternative of the above mentioned diagnostic methods, loop-mediated isothermal amplification (LAMP) was not yet available for ToBRFV. This technique was originally designed by Notomi et al. [15], in which six specific primers named internal primers (F2-F1c/B2-B1c), external primers (F3/B3) and loop-specific primers (LoopF/LoopB) were used. Compared to other methods, LAMP has advantages due to its rapidity, specificity and simplicity. For example, a typical LAMP reaction can be completed within 30 min, and it is highly specific as it can detect six gene regions of the target sequence by the six primers. Furthermore, LAMP does not need specialized equipment, such as a thermocycler, but can be conducted in warm water bath, thermal blocks, or oven, and does not require electrophoresis for detection and identification of the reaction product(s).

LAMP diagnostic protocols have been developed for many plant diseases [16–20]. However, there was no such protocol available for ToBRFV. The purpose of the present study was to develop a LAMP test for detection of ToBRFV and compare it with PCR and qPCR tests. Our results indicated that the LAMP method was sensitive and specific with potential to be developed into a field-friendly diagnostic test.

## Materials and Methods

### Chemicals and standard techniques

All chemicals and equipment were purchased from Fisher Scientific Canada (Ottawa, ON) unless otherwise specified. Extraction of genomic DNA and total RNA from plant samples was conducted using a DNeasy plant mini kit (Qiagen Canada, Toronto, ON) and a RNeasy plant mini kit (Qiagen Canada), respectively. Synthesis of the first-strand cDNA from the extracted total RNA was conducted using a QuantaBio qScript cDNA synthesis kit (VWR Canada, Edmonton, AB). Primers and gBlocks were synthesized by Integrated DNA Technologies (Coralville, IA). LAMP, polymerase chain reaction (PCR) and reverse transcriptase (rt)-PCR were conducted in a Proflex 96-well PCR cycler. Quantitative (q) PCR was performed in a CFX96 touch real-time PCR detection system (Bio-Rad Canada, Mississauga, ON).

### LAMP primer design and pre-selection

A query of the national center for biotechnology information (NCBI) whole genome sequence (WGS) database (https://www.ncbi.nlm.nih.gov/genome) with the term “tomato brown rugose fruit virus” resulted in one entry (tested on Feb 25, 2020) with an International Nucleotide Sequence Database Collaboration (INSDC) number KT383474. Based on this WGS, ten sets of LAMP primers were designed using LAMP Designer (http://www.premierbiosoft.com/isothermal/lamp.html), an online tool designing LAMP primers, and screening them for specificity. Using the WGS of KT383474 as query, a BLASTn was conducted against the NCBI database. From the result of the BLAST, all available ToBRFV WGS (seven in total as confirmed by Feb 25, 2020), three selected TMV WGS and two selected ToMV WGS were retrieved. Alignment of these 12 sequences was conducted using Clustal Omega (https://www.ebi.ac.uk/Tools/msa/clustalo). The output of the alignment was transferred into a Microsoft Word file and the file was named Align.doc. Based on Align.doc, the specificity of all primers in the ten primer sets was manually checked. If the last five nucleotides at the 3’ end of a primer matched with any of the non-targets (the three TMV and the two ToMV) or did not match with one of the targets (the seven ToBRFV), the entire primer set that this primer belongs to was discarded. After this pre-selection, three LAMP primer sets remained and one set was selected for use in the subsequent studies. The selected primer set was hereafter referred as LAMP primers and their locations and sequences are given in Fig. 1 and Table 1, respectively.

**Table 1.**
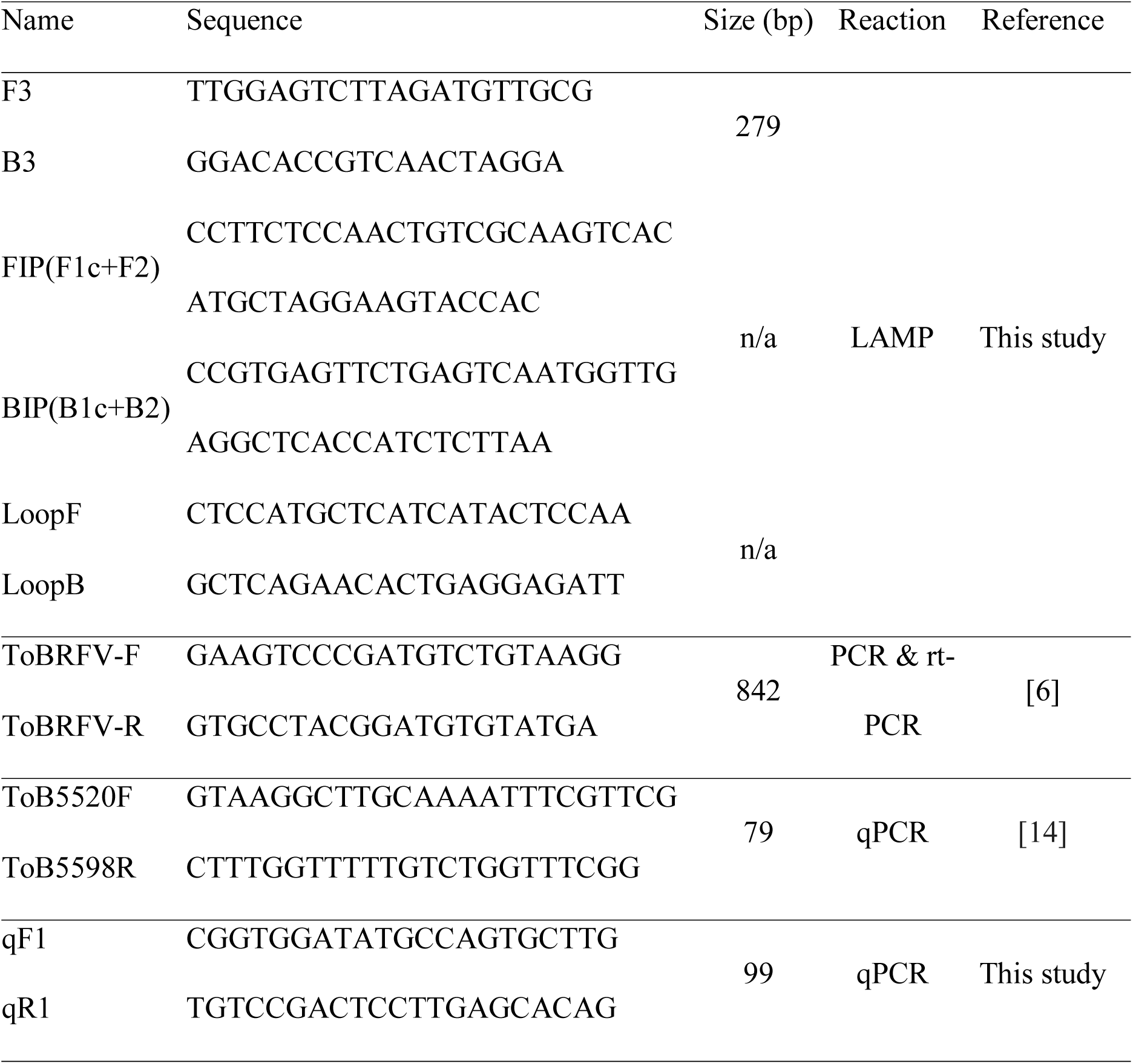
Primers used in this study

**Fig. 1.**
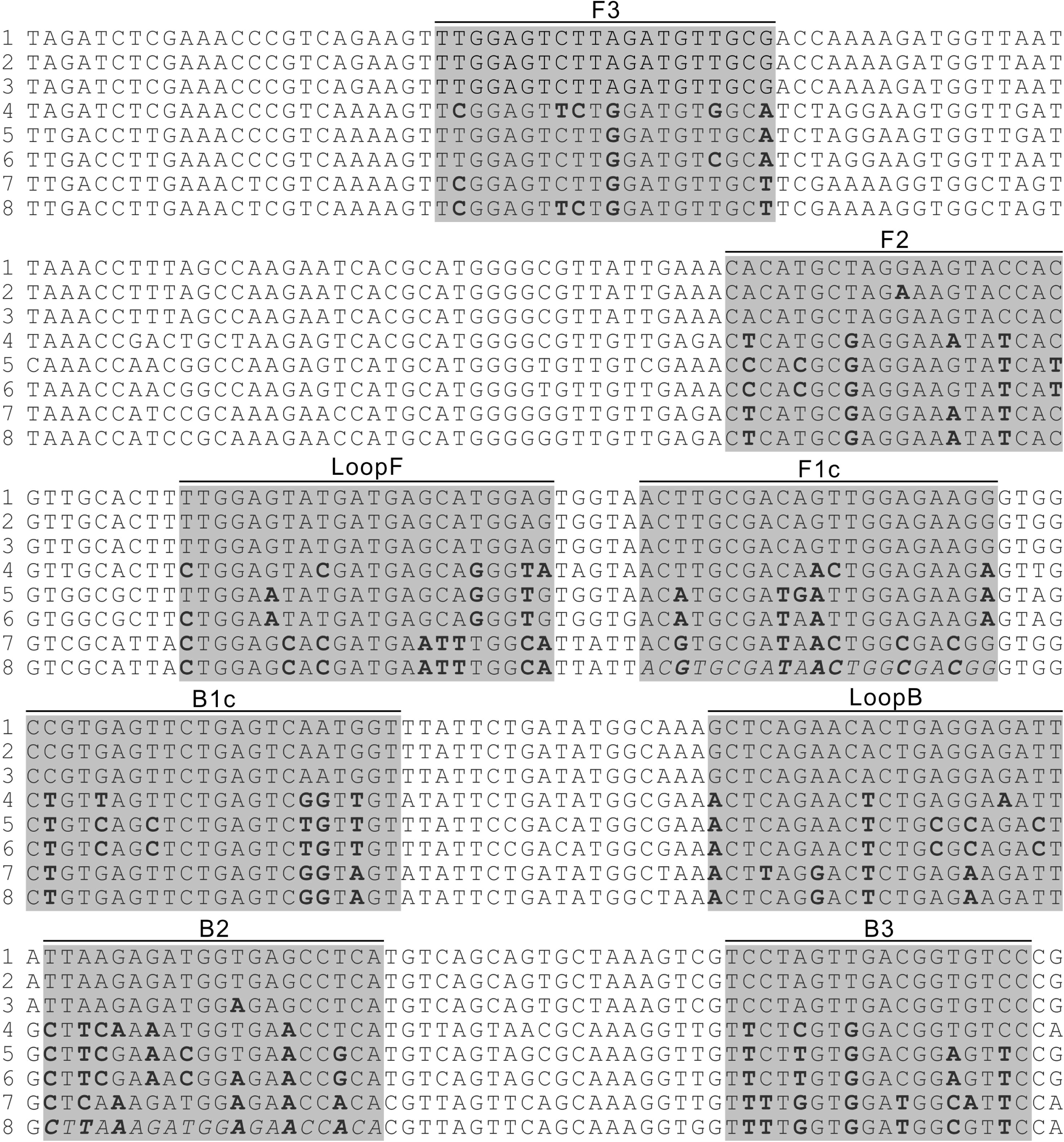
Alignment of partial genomic sequences of selected ToBRFV, TMV and ToMV strains. Locations of LAMP primers were highlighted and the primer name indicated above. Degenerated nucleotides within the primer binding sites were bolded. Italic letters indicate the sequences present in gBlock^ΔF1c^ and gBlock^ΔB2^ that differentiate the two gBlocks from gBlock^ToBRFV^. Line 1, represents five ToRBFV strains. Lines 2 and 3, are sequences from two ToBRFV strains. Lines 4-6 are sequences from three TMV strains. Lines 7 and 8 are sequences from two ToMV strains. (Detailed description of the virus strains is list in Table S1).

### Design of gBlocks and qPCR primers

In the file Align.doc, one genomic region of 999 nt containing all binding sites of the LAMP primers was selected. Three 999-nt double-stranded DNA fragments, corresponding to the RNA sequence of this genomic region in one selected ToBRFV strain (INSDC no: KT383474; nt 1657-2655), one selected TMV strain (INSDC no: FR878069; nt 1649-2647) and one selected ToMV strain (INSDC no: MH507166; nt 1653-2651) were synthesized as gBlocks. These gBlocks were named gBlock^ToBRFV^, gBlock^TMV^ and gBlock^ToMV^, respectively. In addition, two gBlocks, in which the sequences of the binding site of primer F1c or B2 on gBlock^ToBRFV^ were replaced with the corresponding sequences in one of the ToMV strains (DQ873692), were synthesized and named gBlock^ΔF1c^ and gBlock^ΔB2^, respectively (Table 1 and Fig. 1). Based on the sequence of gBlock^ToBRFV^, a pair of qPCR primers was designed using the Primer3 software (http://primer3.ut.ee). This primer pair was name qF1/qR1 and their sequences are given in Table 1.

### LAMP assays

All LAMP reactions were conducted in the WarmStart colorimetric LAMP master mix (NEB Canada, Whitby, ON). Each reaction was 25 μL in total and contained 1 μL template. Quantities of other components in each reaction followed the NEB’s instructions for the master mix. All reactions were conducted in individual 200-μL PCR tubes. The reaction program consisted of only one step in which the tubes were incubated at 65°C for 30 minutes. After the incubation, the reactions were checked visually and the results were recorded by photography.

### PCR, rt-PCR and qPCR

Each reaction of PCR, rt-PCR and qPCR was 25 μL containing 1 μL template and 0.25 μM of each primer. PCR and qPCR was conducted in Promega PCR master mix and SsoAdvanced universal SYBR green supermix (Bio-Rad Canada), respectively. The PCR program consisted of an initial denaturation at 94°C for 3 min, followed by 40 cycles of denaturation at 94°C for 30 s, annealing at 55°C for 45 s and extension at 72°C for 1 min, and a final extension at 72°C for 5 min. The qPCR program consisted of an initial denaturation step of 95°C for 2 min, followed by 40 cycles of 5 s at 95°C and 1 min at 60°C. After completion of the qPCR amplification, a melting curve analysis was run to evaluate the amplification specificity. rt-PCR was conducted with a OneStep RT-PCR kit (Qiagen Canada) by following the manufacturer’s instructions.

### Assays on gBlocks

Each gBlock was dissolved in water and the concentration was adjusted to 100 fM, which roughly equals to 60000 molecules/μL. From this concentration a set of 10× serial dilutions were prepared for each gBlock down to a concentration of 0.06 molecules/μL (seven solutions for each gBlock). Using the serial dilutions of all gBlocks as templates, LAMP was conducted. Using the serial dilutions of gBlock-ToBRFV as templates, PCR and qPCR were conducted with primers ToBRFV-F/ToBRFV-R [6](Ling et al. 2019) and qF1/qR1 (Table 1), respectively. In all assays, sterile, DNA-free water was used as the negative control. All LAMP, PCR and qPCR were repeated using the same preparations of serial dilutions.

### Assays on genomic DNA from health plants

Genomic DNA was extracted from young leaves of Tomato (var. Cherry Nebula), pepper (var. Tabasco), potato (*Solanum tuberosum* var. Norland) and tobacco (*Nicotiana tabacum* unknown variety). Using 40 ng of genomic DNAs as templates, LAMP was conducted individually for each of the four host plants. The LAMP assay was repeated on alternative DNA preparations.

### Assays on RNA of infected plants

Total RNA was extracted from tomato plants with symptoms of ToBRFV infection. Five RNA samples, confirmed to be ToBRFV positive, were provided to APHL by the Plant Disease Clinic of University of Guelph and named ‘original RNA samples’. The APHL tested the five original RNA samples again with rt-PCR using the primer pair ToBRFV-F/ToBRFV-R, and with LAMP using the designed LAMP primers. In addition, a 2-μL subsample was taken from each of the five original RNA samples; the five subsamples were pooled and diluted 30× to form a 300-μL RNA mixture. From this mixture, a set of eight 10-fold dilution series was prepared. rt-PCR and LAMP were conducted on all eight dilutions using the above indicated primers. In all assays, sterile, RNA-free water was used as the negative control. All assays were repeated using the same RNA preparations.

### Assays on cDNA derived from the infected plants

A 2-μL subsample was taken from each of the five original RNA samples; the five subsamples were pooled, from which the first-strand cDNA was synthesized and the final cDNA solution was adjusted to 300 μL. From this cDNA solution, a set of eight 10-fold dilution series was prepared. These eight dilutions were used as template for PCR, LAMP and qPCR reactions. The primer pair ToBRFV-F/ToBRFV-R was used in PCR. Primer pairs ToB5520F/ToB5598R and qF1/qF2 were used in qPCR. In all assays, sterile DNA-free water was used as the negative control. All assays were repeated using the same cDNA preparations.

### Statistics

In all qPCR assays, each sample was tested with three technical repeats. The average and standard deviation of the quantification cycle (Cq) values from the three repeats were calculated using MS Excel. Calculation of Poisson distribution was conducted with an online Poisson distribution calculator (https://stattrek.com/online-calcμLator/poisson.aspx).

## Results

### LAMP on the gBlock

In all LAMP reactions, changing color of the reaction mixture from pink to yellow indicated a positive result, which was hereafter referred to as positive signal. With the designed LAMP primers, no positive signal could be generated from gBlock^TMV^ and gBlock^ToMV^ (Fig. 2A). Positive signals were generated from gBlock^ToBRFV^ when its molecule number was more than or equal to 6 per 25-μL reaction (Fig. 2A). When gBlock^ΔF1c^ and gBlock^ΔB2^ were used as the templates, positive signals could be generated when their molecule numbers were more than or equal to 600 per 25-μL reaction; there was no positive signal when the molecule numbers were less than or equal to 60 per 25-μL reaction (Fig. 2B). These results confirmed that the LAMP primers were ToBRFV-specific. When the 6 molecules/μL gBlock^ToBRFV^ sample was repeatedly tested by LAMP, 35 out of 40 reactions were found to be positive (Fig. 2C). This result indicated that the LAMP assay could detect as little as 6 molecules in a 25-μL reaction more than 87% of the time. In contrast, PCR could detect the target when the molecule number of gBlock^ToBRFV^ was more than or equal to 600 per 25-μL reaction (Fig. 2D). In qPCR assay, Cq values were generated from all of the three repeated reactions when the molecule number of gBlock ToBRFV was more than or equal to 6 per 25 μL; when the molecule number was 0.6 per 25 μL, only one out of the three reactions generated a Cq value (Fig. 2E). These results indicated that the sensitivity of the LAMP assay on the gBlock was similar to that of qPCR but 100 times higher than that of PCR. Repeated assays of LAMP, PCR and qPCR produced results similar to Figs. 2A and 2C, Fig. 2D and Fig. 2E, respectively.

**Fig. 2.**
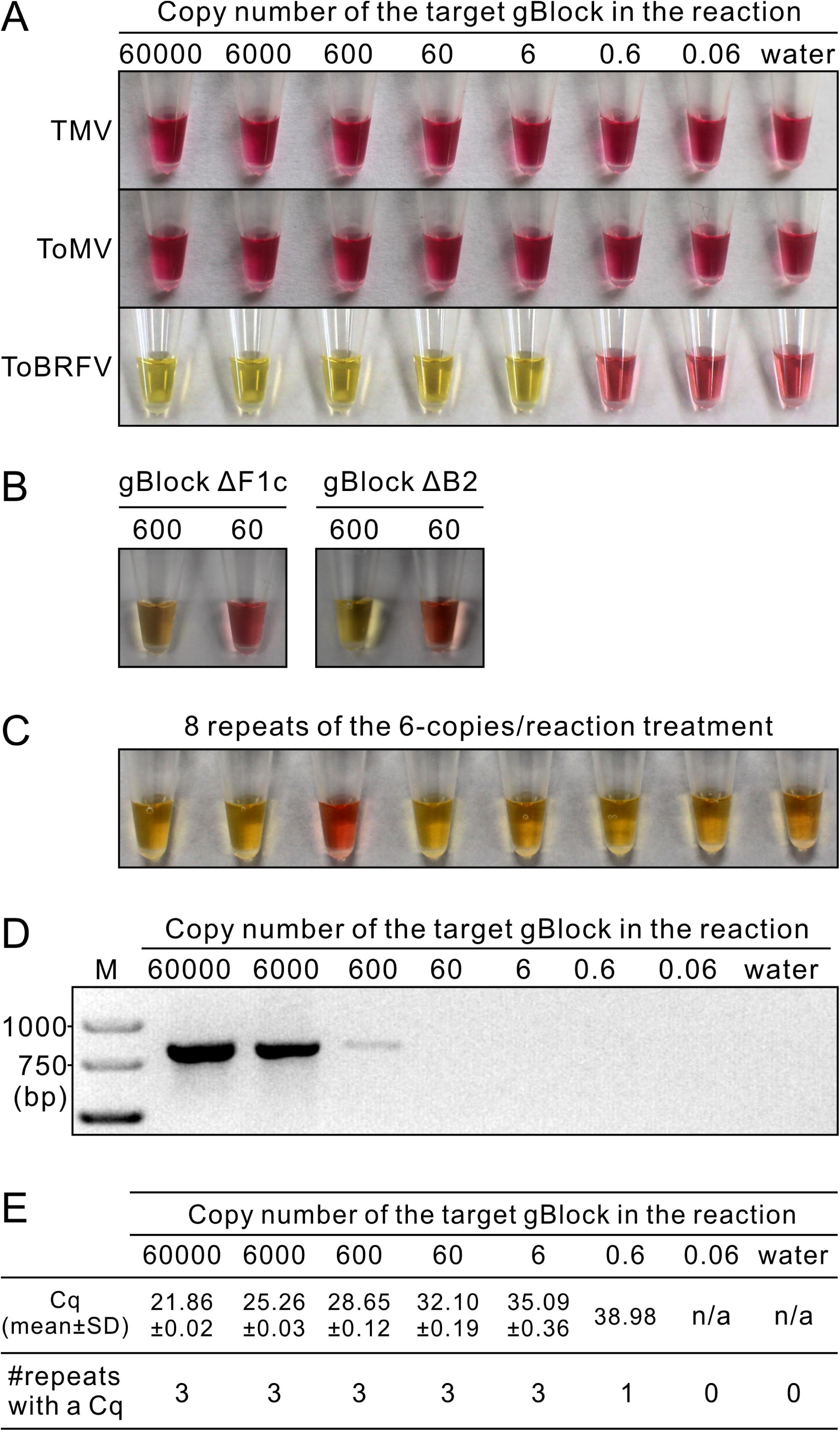
LAMP (A-C), PCR (D) and qPCR (E) tests on gBlocks. A, LAMP test on serial dilutions of gBlock^TMV^, gBlock^ToMV^ and gBlock^ToBRFV^. B, LAMP test on two dilutions of gBlock^ΔF1c^ and gBlock^ΔB2^. C, Repeated LAMP tests on gBlock^ToBRFV^ at the concentration of 6 molecules per reaction. 40 repeats were conducted but only eight were shown; 35 out of the 40 were positive. D, PCR test on serial dilutions of gBlock^ToBRFV^ using the primer pair ToBRFV-F/ToBRFV-R. M, GeneRuler Express DNA Ladder. E, qPCR test on serial dilutions of gBlock^ToBRFV^ using the primer pair qF1/qR1.

### LAMP on plant genomic DNA

When genomic DNA from healthy plants of four *Solanaceae* species was used as the template in LAMP, no positive signal could be generated (data not shown). This result confirmed the specificity of the LAMP primers.

### LAMP on RNA samples

When RNA was used as the template, LAMP generated positive signals from all five original RNA samples (Fig. 3A). This result was confirmed by rt-PCR (Fig. 3B). When the serial dilutions were used as the templates, LAMP could generate positive signals from the dilutions 1-7 (Fig. 3C); in contrast, rt-PCR could only produce a band from dilutions 1-5 (Fig. 3D). This result indicated that LAMP was 100 times more sensitive than rt-PCR on RNA samples. Repeated assays of LAMP and PCR produced results similar to Figs. 3A and 3C and Figs. 3B and 3D, respectively.

**Fig. 3.**
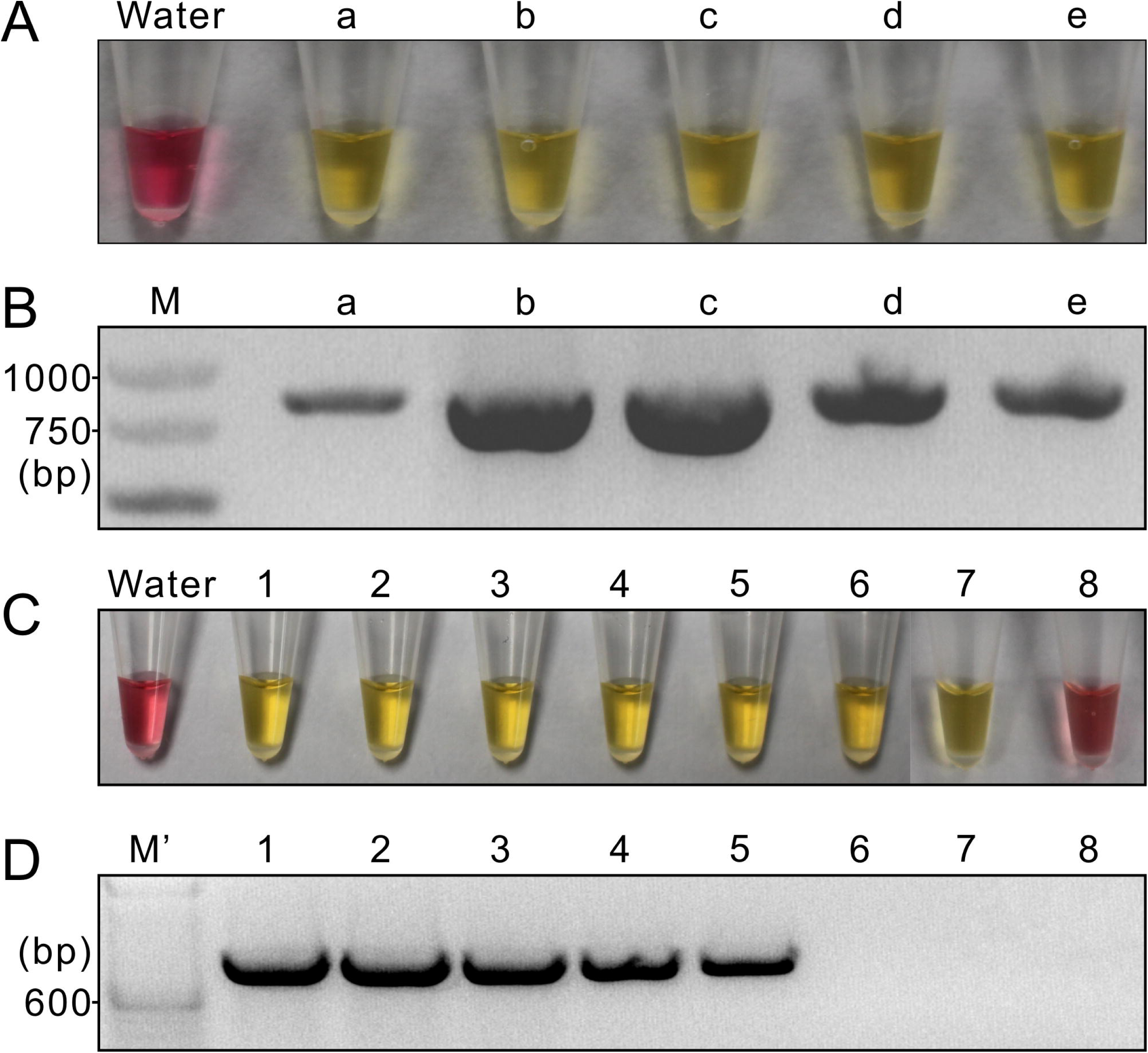
LAMP (A and C) and rt-PCR (B and D) tests on total RNA extracted from five tomato samples infected by ToBRFV. A and B, Tests on five original RNA samples. a, from tomato fruit; b, from tomato leaf; c-e, from tomato flesh. M, GeneRuler express DNA ladder. C and D, Tests on serial dilutions (1-8) of a RNA sample pooled from same volumes of each of the five original RNA samples. M’, TrackIt 100 bp DNA ladder.

### Efficiency of LAMP on cDNA

When the serial dilutions of cDNA were used as the templates, LAMP generated positive signals from samples 1-6 dilutions (Fig. 4A). In contrast, PCR generated positive bands from samples 1-4 (Fig. 4B). qPCR assays using alternative primer sets showed similar sensitivity as LAMP (Fig. 4C). Repeated assays of LAMP, PCR and qPCR produced results similar to Fig. 4A, Fig. 4B and Fig. 4C, respectively. These results indicated that the sensitivity of the LAMP assay on the cDNA samples was similar to that of qPCR but 100 times higher than that of PCR.

**Fig. 4.**
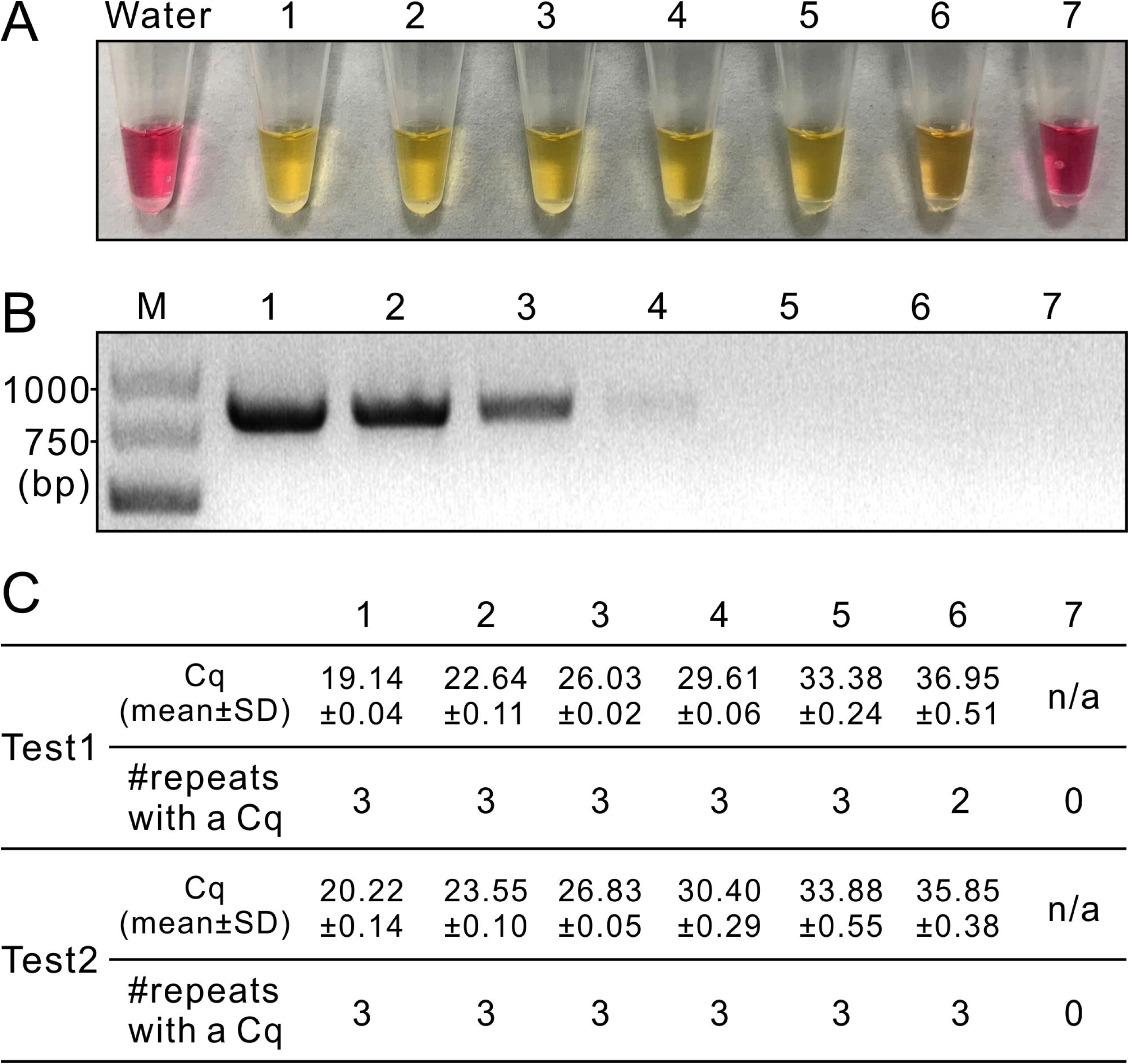
LAMP (A), PCR (B) and qPCR (C) tests on serial dilutions (1-7) of cDNA synthesized from a RNA sample pooled from same volumes of each of the five original RNA samples. M, GeneRuler express DNA ladder. Primer pair used in PCR was ToBRFV-F/ToBRFV-R and those used in qPCR were qF1/qR1 for Test 1 and ToB5520F/ToB5598R for Test 2.

## Discussion

### Specificity of the LAMP

LAMP is more specific than other PCR-based techniques because LAMP has six specific primers. In this study, all LAMP primers were designed to be specific to all ToBRFV strains with an available WGS, and non-specific to TMV and ToMV. TMV and ToMV are closely related to ToBRFV [3], which is supported by the fact that WGS of TMV and ToMV were always the top BLAST hits when the NCBI database was queried with the ToBRFV WGS. When the LAMP primers were tested on the gBlock^TMV^ and gBlock^ToMV^, as well as genomic DNA from various *Solanaceae* plants, no signal could be generated, further confirming the specificity of the LAMP primers to ToBRFV. Moreover, when the LAMP primers were tested on gBlock^ΔF1c^ and gBlock^ΔB2^, which differed from gBlock^ToBRFV^ by only on a few nt on the binding sites of primer F1c and B2, respectively, no positive signal could be generated at concentrations of ≤ 60 molecules/25μL. This result supported the conclusion that the LAMP primers were highly specific to ToBRFV.

### Sensitivity of the LAMP

On the serial dilutions of the gBlocks or cDNA, LAMP was 100 times more sensitive than rt-PCR and similar in sensitivity as qPCR. As well, on the serial dilutions of the RNA samples, LAMP was 100 times more sensitive than rt-PCR. These results were comparable with those of LAMP testing on other RNA viruses. For examples, LAMP was 100 times more sensitive than rt-PCR on the detection of *Citrus leaf blotch virus* [21] and *Little cherry virus 1* [22]; on *Potato virus X* [23] and *Sugarcane streak mosaic virus* [24], LAMP was 10 times more sensitive than rt-PCR. Compared to rt-qPCR, LAMP was similarly sensitive for detection of *Onion yellow dwarf virus* [25] and *Potato leafroll virus* [26].

Testing primers on gBlocks allows more accurate calculation of primer efficiency. In this study, we obtained data that 35 out of 40 LAMP reactions generated positive signals when the template gBlock concentration was 6 molecules/μL. At a very low concentration, the distribution of DNA particles in aliquots follows a Poisson distribution [27]. Using the Poisson distribution calculator, we could calculate that the possibility of obtaining ≥ 4 gBlock molecules in a 1-μL aliquot from a 6 molecules/μL solution was 85%, which is comparable to the data from the LAMP test (35/40 = 88%). Since gBlock is double-stranded and the virus RNA is single-stranded, we concluded that the LAMP could detect the virus RNA in a concentration of as low as 8 viruses per reaction. This is comparable to qPCR results obtained in this study and to the theoretical limit of qPCR detection (3 molecules) proposed by Forootan et al. [28. The estimated efficiency of the RNA extraction kit is approximately 40 to 80% (the ratio between the molecule numbers of extracted genomic RNA and the number of virus particles in the sample) when the sample was 100-mg tomato leaf tissue and the final RNA volume was 50 μL (Feng, unpublished data). Since 1 μL template was used in the LAMP test, we concluded that the LAMP test could detect ToBRFV when a 100-mg plant sample contains as little as 1,000 virus particles (8÷40%×50). While this level of detection is encouraging, it is important to note that the minimal number of virus particles necessary to cause plant infection could be very low. For example, one Tobacco etch virus (TEV) particle is sufficient to initiate a systemic infection [29]. We want to emphasize here that more improvement is required on the commonly used diagnostic protocols, and that the efficiency of a molecular-based diagnostic protocol is determined not only by the efficiency of the detection technique only, but also by other factors such as the efficiency of DNA/RNA extraction.

### Time-saving and simplicity of LAMP

Compared to PCR, rt-PCR or rt-qPCR, LAMP is time saving. For example, in the present study, testing one DNA, RNA or cDNA sample by LAMP could be completed within 35 min. Testing eight samples could be completed in less than one hour. In contrast, testing one or eight samples by PCR or rt-PCR needed more than three hours (for PCR and electrophoresis) and testing by rt-qPCR would need more than 2 hours. All LAMP results reported in the present study were generated in a PCR thermocycler. However, identical results were obtained when the LAMP tests were conducted in an oven or a heat block. Thus, for labs where PCR or qPCR thermocycler is not available, LAMP protocol developed in this study is highly recommended for ToBRFV testing.

## Acknowledgements

We thank Dr. Shannon Xuechan Shan from the Plant Disease Clinic of University of Guelph for providing all RNA samples used in this study. This study was supported by a grant from the Canadian Agricultural Partnership (no. 601322) to JF.

## Author Contributions

Conceptualization: Jie Feng.

Data Curation: Jie Feng.

Formal Analysis: Jie Feng.

Funding Acquisition: Jie Feng, Michael W. Harding, David Feindel.

Investigation: Alian Sarkes, Heting Fu, Jie Feng.

Methodology: Jie Feng.

Project Administration: Jie Feng.

Resources: Jie Feng, David Feindel.

Supervision: Jie Feng.

Validation: Jie Feng.

Visualization: Jie Feng.

Writing – Original Draft Preparation: Jie Feng.

Writing – Review & Editing: Jie Feng, Michael W. Harding.

## Notes

#### Summary of Updates

Author Contributions

