## Supplementary material for "Development and evaluation of a loop-mediated isothermal amplification (LAMP) assay for the detection of *Tomato brown rugose fruit virus* (ToBRFV)": Table S1

| **Table S1. Sequences used in the alignment in Fig. 1.** | | | | |
| --- | --- | --- | --- | --- |
| No | Species | GenBank No. | Origin | Nucleotides |
| 1 | ToBRFV | KT383474 | Jordan | 2277-2581 |
| 1 | ToBRFV | MK133095 | Germany | 2274-2578 |
| 1 | ToBRFV | MN167466 | Italy | 2274-2578 |
| 1 | ToBRFV | MN182533 | UK | 2562-2866 |
| 1 | ToBRFV | MK648157 | Jordan | 2271-2575 |
| 2 | ToBRFV | MK133093 | Germany | 2273-2577 |
| 3 | ToBRFV | KX619418 | Israel | 2275-2579 |
| 4 | TMV | FR878069 | USA | 2269-2573 |
| 5 | TMV | MH595921 | China | 2269-2573 |
| 6 | TMV | KY810785 | Slovenia | 2270-2274 |
| 7 | ToMV | MH507166 | Korea | 2272-2576 |
| 8 | ToMV | DQ873692 | Germany | 2272-2576 |
